# Cryo-EM structure of the fully-loaded asymmetric anthrax lethal toxin in its heptameric pre-pore state

**DOI:** 10.1101/2020.04.07.029140

**Authors:** Claudia Antoni, Dennis Quentin, Alexander E. Lang, Klaus Aktories, Christos Gatsogiannis, Stefan Raunser

## Abstract

Anthrax toxin is the major virulence factor secreted by *Bacillus anthracis*, causing high mortality in humans and other mammals. It consists of a membrane translocase, known as protective antigen (PA), that catalyzes the unfolding of its cytotoxic substrates lethal factor (LF) and edema factor (EF), followed by translocation into the host cell. Substrate recruitment to the heptameric PA pre-pore and subsequent translocation, however, are not well understood. Here, we report three high-resolution cryo-EM structures of the fully-loaded anthrax lethal toxin in its heptameric pre-pore state, which differ in the position and conformation of LFs. The structures reveal that three LFs interact with the heptameric PA and upon binding change their conformation to form a continuous chain of head-to-tail interactions. As a result of the underlying symmetry mismatch, one LF binding site in PA remains unoccupied. Whereas one LF directly interacts with a part of PA called α-clamp, the others do not interact with this region, indicating an intermediate state between toxin assembly and translocation. Interestingly, the interaction of the N-terminal domain with the α-clamp correlates with a higher flexibility in the C-terminal domain of the protein. Based on our data, we propose a model for toxin assembly, in which the order of LF binding determines which factor is translocated first.

## Introduction

Anthrax is a life-threatening infectious disease that affects primarily livestock and wild animals, but can also cause high mortality in humans (1). During the early and late steps of infection with the Gram-positive bacterium *B. anthracis*, the tripartite anthrax toxin is secreted as major virulence factor in order to kill host immune cells such as macrophages or neutrophils (2,3). Like other AB-type toxins, it is composed of a surface binding/translocation moiety, the protective antigen (PA, 83 kDa), and two cytotoxic subunits, lethal factor (LF, 90 kDa) and edema factor (EF, 93 kDa) (4,5).

To execute their toxicity, both the zinc-dependent metalloproteinase LF and/or the adenylate cyclase EF need to enter the host cytoplasm (6,7). For that purpose, PA monomers first attach to the cell surface through binding to one of the two known membrane receptors, capillary morphogenesis gene 2 (CMG-2) and tumor endothelial marker 8 (TEM8) (8,9). After cleavage by furin-like proteases, the truncated 63 kDa-sized PA monomer oligomerizes either into homo-heptamers (PA_7_) or homo-octamers (PA_8_) (10-12). These ring-shaped oligomers, enriched in lipid raft regions, are in a pre-pore conformation as they do not penetrate the host membrane (13). Due to the enhanced stability of PA_8_ under diverse physiological conditions, it is proposed that the octameric form could circulate in the blood to reach and exert toxicity even in distant tissues (14). This suggests that both oligomeric forms play an important role in intoxication, endowing *B. anthracis* with greater versatility against its host.

In the next step, the holotoxin is assembled by recruiting LFs/EFs. While PA_8_ can bind up to four factors, only three of them can simultaneously bind to PA_7_. Both enzymatic substrates bind to the upper rim of the PA oligomer via their N-terminal domains in a competitive manner (15). Loaded complexes are then endocytosed (16,17), followed by a conformational change from the pre-pore to pore state which is triggered by the low pH in the endosome (18). The central feature of the pore state is an 18 nm long 14-stranded β-barrel that spans the endosomal membrane with its narrowest point in the channel lumen being ∼6 Å in width (19). To pass through this hydrophobic restriction, called Φ-clamp, the substrate needs to be unfolded prior to translocation (20).

Structural and functional studies on the pre-pore PA octamer bound to four LFs revealed that an amphipathic cleft between two adjacent PA protomers, termed α-clamp by Krantz and coworkers, assist in the unfolding process (21). More specifically, the first α-helix and β-strand (α1-β1) of LF almost completely unfold and change their position respective to the rest of the protein when interacting with the α-clamp of the PA oligomer (21). After transition into the pore state, the unidirectional translocation of LF is driven by a proton-motive force, comprising the proton gradient between the two compartments and the membrane potential. It is thought that the acidic pH present in the endosome destabilizes the LF and thus promotes unfolding of its N-terminus (22). Ultimately, it is believed that the translocation follows a ‘charge-dependent Brownian ratchet’ mechanism (23). The required unfolding and refolding of translocated enzymes is facilitated *in vivo* by chaperones, but can occur *in vitro* without the need of accessory proteins (24,25).

Crystallographic studies provided us with structural insights pertinent to the molecular action of the anthrax toxin. This includes structures of the individual complex subunits such as LF, EF and the PA pre-pore in both, its heptameric and octameric form (12,26-29). The PA monomer was also co-crystallized with its receptor CMG-2, delineating the surface attachment to the host cell in molecular detail. More recently, the elusive pore state of PA_7_ was determined by electron cryo-microscopy (cryo-EM) in which Jiang *et al*. made use of an elegant on-grid pore induction approach (30).

In contrast, high resolution information on holotoxin complexes is rather scarce. The only obtained crystallographic structure is the aforementioned PA_8_ pre-pore in complex with four LFs (21). In this structure, however, the C-terminal domain of LF is absent. Unlike PA_8_, loaded PA_7_ was mainly studied by cryo-EM (31-35), presumably because its asymmetry impeded crystallization efforts. Earlier this year, the PA_7_ pore state decorated with a single LF molecule and with up to two EF molecules was determined, in which it was shown that EF undergoes a large conformational rearrangement as opposed to LF (36). However, cryo-EM studies of the loaded heptameric pre-pore were so far limited to a resolution of ∼16 Å (31,32,34). In addition, the number of LFs bound to PA_7_ varied between one and three in these structures.

Here, we present three cryo-EM structures of the fully loaded anthrax lethal toxin in the heptameric pre-pore state (PA_7_LF_3_), in which three LF molecules are bound to the rim of the PA_7_ ring, forming a continuous chain of head-to-tail interactions. The position and conformation of the LFs, however, varies between the structures. Unexpectedly, only one of three LFs interacts with the α-clamp of PA, adopting the “open” conformation as reported in the PA_8_LF_4_ structure (21). Since we could neither observe a similar interaction for the other two LFs, nor them being in the “closed” conformation, we propose that they adopt an “intermediate” state between holotoxin assembly and translocation. Our findings allow us to propose a model for anthrax lethal toxin assembly, in which the LF translocation sequence is dictated by the order of LF binding.

## Results

### Structure of the fully-loaded anthrax lethal toxin in the heptameric pre-pore state

To ensure that our purified and reconstituted PA_7_ complexes (Materials and Methods) are indeed intact, we tested their membrane insertion capacity by reconstituting them in liposomes or nanodiscs (Fig. S1). We then evaluated different molar ratios of LF:PA_7_ and only obtained fully-loaded anthrax lethal toxin (PA_7_LF_3_) when using a 10:1 molar ratio as judged by size exclusion chromatography (Fig. S2A, B).

We then determined the structure of the PA_7_LF_3_ pre-pore complex by single particle cryo-EM to an average resolution of 3.5 Å. However, the densities corresponding to LF represented a mixture of assemblies and were partly unassignable (Fig. S3). This can be either due to the symmetry mismatch that emerges when three lethal factors bind simultaneously to PA_7_ or to possible different conformations of the individual LFs bound to PA_7_. To address these points, we established an image processing workflow that includes sequential 3-D classifications and rotation of classes (Fig. S3). This resulted in three reconstructions with resolutions of 3.8 Å, 4.2 Å and 4.3 Å that differed in the position of the third LF bound to PA_7_ (Fig. 1A, B, Fig. S2, S3, S4) and the conformation of LF (Fig. 1C, Fig. S2, S3, S4). The densities corresponding to the lethal factors in the 4.3 Å structure were not resolved well enough to allow the fitting of an atomic model (Fig. S2F, I, S4B). Therefore, we proceeded with the remaining two structures, combined the two particle stacks and masked out the density of the third LF to improve the resolution of the rest of the complex to 3.5 Å (Fig. S2H, S3, S4D). Using a combination of the maps, we then build atomic models for the 3.8 Å and 4.2 Å reconstructions (Fig. 1A, B, Table S1).

**Figure 1.**
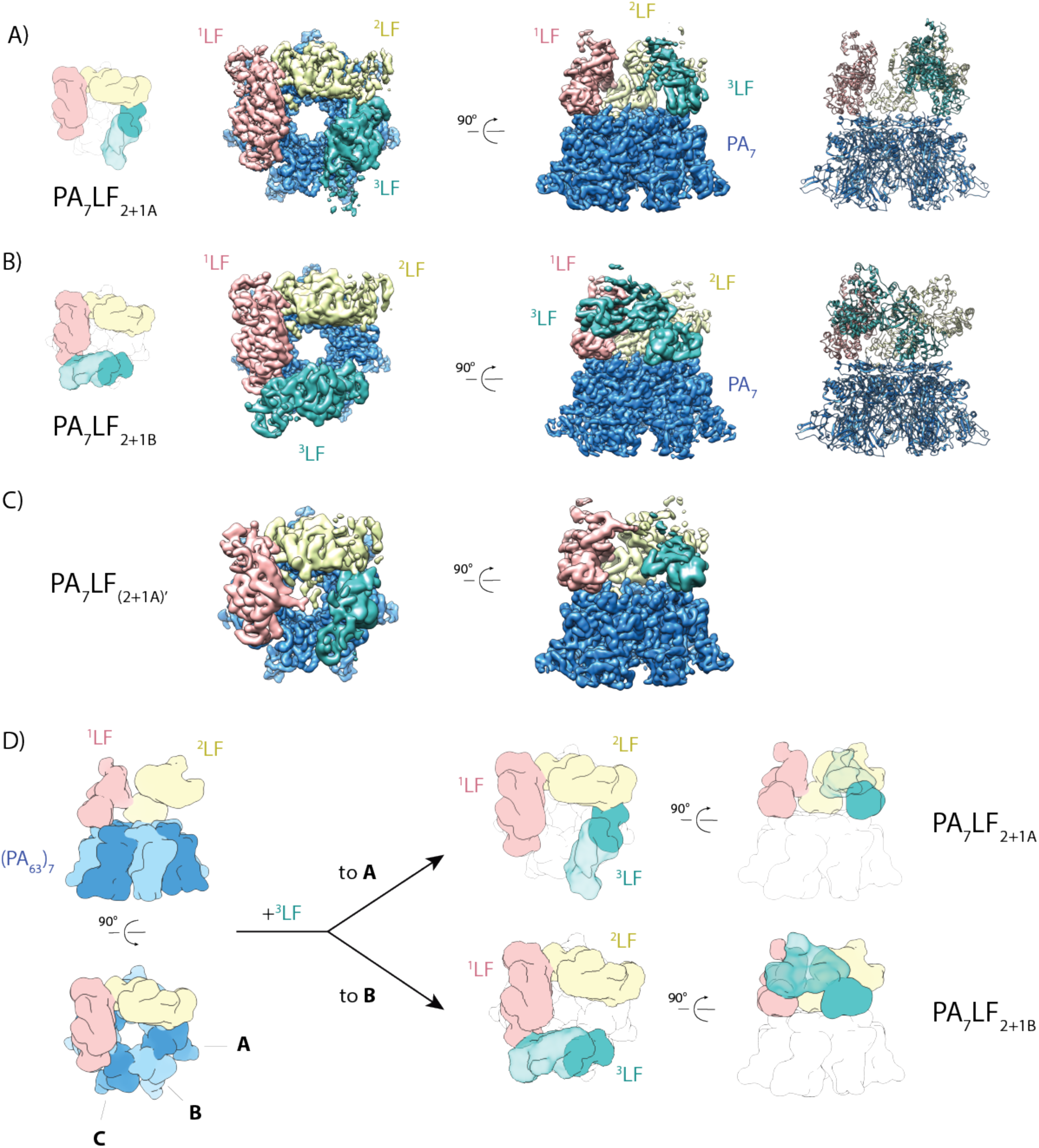
Cryo-EM structures of the PA_7_LF_3_ complexes. (**A**) Top view and side view of the color-coded segmented cryo-EM density map of PA_7_LF_2+1A_, with PA_7_ in blue, ^1^LF in pink, ^2^LF in gold and ^3^LF in cyan. Three lethal factors bind to the PA_7_ ring and form a continuous chain of head-to-tail interactions. Schematic representation is shown on the left, corresponding atomic model on the right. (**B**) Same as in (A) for the PA_7_LF_2+1B_ complex. (**C**) Same as in (A) for the PA_7_LF_(2+1A)’_ complex. Notably, two LFs interact in their peripheral region (C-terminal domain) with each other close to the central axis. Segmented maps are shown at different thresholds for visualization. (**D**) Schematic representation of the last step in PA_7_LF_3_ toxin assembly, in which the third lethal factor can bind to one of two empty PA sites, resulting in two different complexes, PA_7_LF_2+1A_ and PA_7_LF_2+1B_. Top and side views are shown, with the same color code as in (A), except that PA protomers alternate in light and dark blue.

The structures reveal that PA_7_ forms a seven-fold symmetric ring structure with a ∼25 Å wide central opening. With the exception of a few poorly resolved loop regions in the periphery of PA_7_, our structures almost perfectly superimpose with the crystal structure of the PA_7_ pre-pore (PDB:1TZO; RMSD of 0.92 Å) (26) (Fig. S5A), indicating that the binding of LF does not induce conformational changes in PA_7_. This is in contrast to Ren et al. who suggested that LF binding results in a distortion of the symmetric PA_7_ ring, thereby facilitating the passage of cargo through the enlarged lumen (31,37). Noteworthy, the 2β2-2β3 loop region (residues 300-323) which is implicated in pore formation was not resolved in our map. This indicates a high flexibility of this loop, which is in line with previous MD simulations (38).

In all PA_7_LF_3_ structures, the three LFs sit on top of PA_7_. The densities corresponding to the LFs show a resolution gradient from the central N-terminal domain which is resolved best to the peripheral C-terminal domain (Fig. S4A-D). This indicates that this region is quite flexible compared to the rest of the toxin complex. The LFs do not only interact with PA_7_ but also form a continuous chain of head-to-tail interactions with each other. Binding of LF to a single PA protomer is mediated via the N-terminal domain of LF, orienting its bulky C-terminal domain such that the adjacent PA protomer is not accessible for binding. In this way a single lethal factor *de facto* occupies two of the seven binding sites of PA_7_. In the chain of LFs, the C-terminal domain of the anterior LF binds to the N-terminal domain of the following one, creating a directionality in the complex (Fig. 1D). Consequently, if two LFs are bound, three free PA binding sites are available, of which only two can potentially be occupied due to steric clashes (Fig. 1D). This results in the two complexes PA_7_LF_2+1A_ and PA_7_LF_2+1B_, that differ in the binding position of the third LF (Fig. 1). Since each LF occupies two potential binding sites in these structures, this leads to a symmetry mismatch and leaves one PA unoccupied.

### Crucial Interactions in the PA_7_LF_3_ complex

LF and PA interact mainly via a large planar interface at which domain I of LF interacts with the LF/EF binding sites of two adjacent PAs (Fig. 2, Fig. S5B). The LF-PA interface is well resolved for all LFs and almost identical in the different structures (Fig. 2A, Fig. S5A, C). The interaction is primarily mediated by an extensive hydrophobic core that is further surrounded by electrostatic interactions (Fig. 2A). The interface in our structure is very similar to the one previously described for PA_8_LF_4_ (21). There, the second LF-PA interface is formed by the N-terminal α-helix of LF that interacts with the α-clamp located at the interface of two PAs. This “open” conformation differs from the “closed” conformation of this region as observed in the structure of the unbound LF (27). When comparing the LFs in our structure with that of the unbound LF, we observed that the C-terminal domain of the LFs in PA_7_LF_3_ is rotated in relation to the N-terminal domain, bringing them closer together (Fig. 3, Movie S1). However, we only found that the N-terminal region of one LF (^2^LF) resides in the α-clamp, adopting the “open” conformation as described for PA_8_LF_4_ (21). In the other LFs (^1^LF, ^3^LF), this region is flexible and not interacting with the α-clamp (Fig. 2B). A steric clash between the loop region (residues 576-579) of ^1^LF and α1-β1 of ^2^LF (Fig. 4A) prevents the N-terminal α-helix of LF from remaining in the “closed” conformation. Since these LFs neither take the “open”, nor the “closed” conformation, we propose that they reside in an “intermediate” conformational state between toxin assembly and translocation. We further hypothesize that ^2^LF is the first of the three lethal factors that is unfolded by PA_7_ and is also the first one to be translocated.

**Figure 2.**
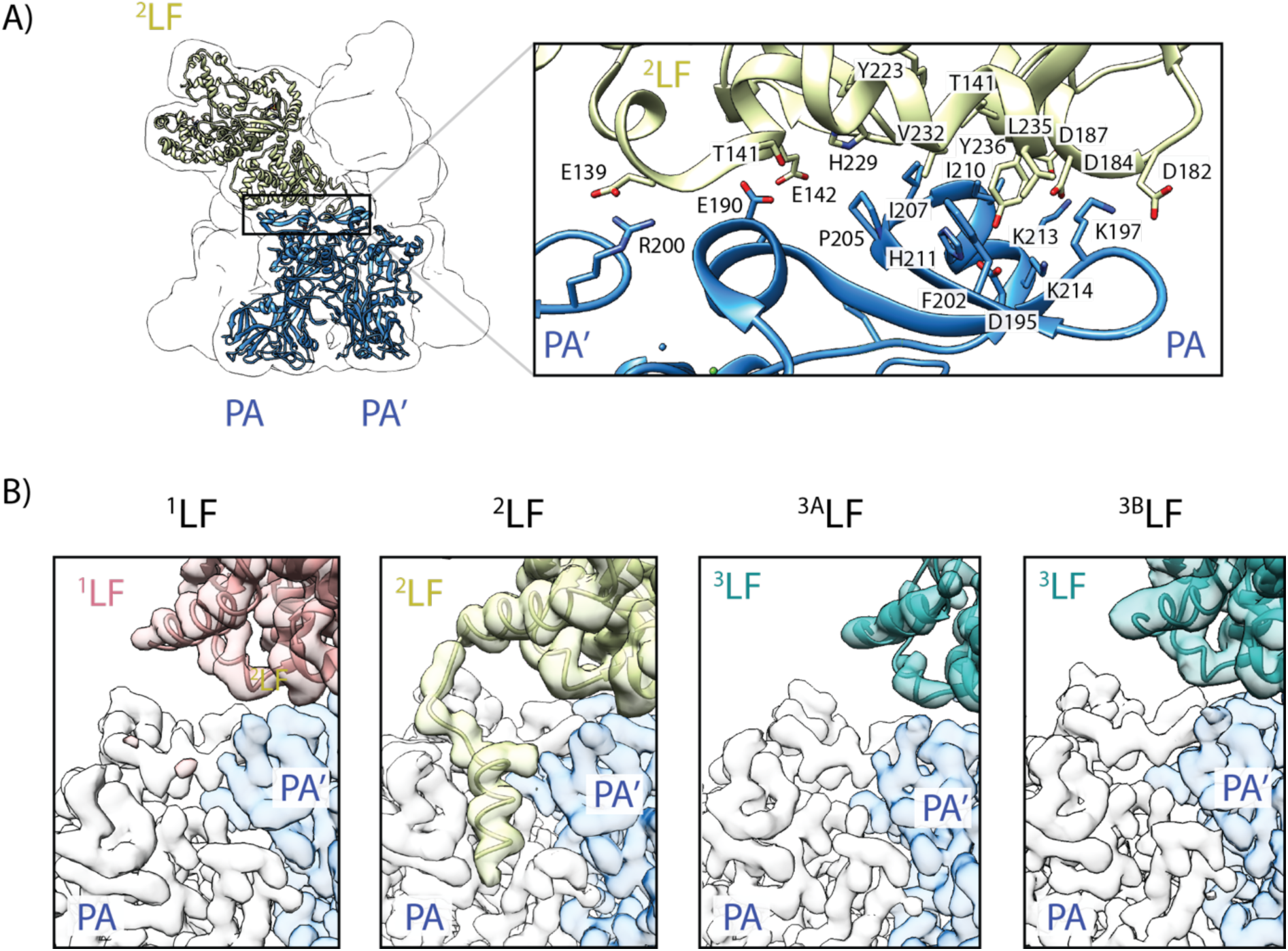
Interfaces between lethal factor and protective antigen. (**A**) The N-terminal domain of LF mediates binding to two adjacent PA molecules, forming a large planar interface. The positions of ^2^LF (gold), PA and PA’(blue) are shown relative to the overall shape of the complex that is represented as transparent, low-pass filtered volume. A black square indicates the interaction interface between all three molecules. The inset shows a close-up of the interaction regions, with contributing residues labeled. They form a central hydrophobic core, that is surrounded by electrostatic interactions. (**B**) The second LF-PA interface is formed by the N-terminal α-helix of LF, interacting with the α-clamp region, located between two adjacent PA molecules. The four panels depict a close-up of this region for the three different LFs (^3^LF can adopt two different positions, i.e. the PA_7_LF_2+1A_ or PA_7_LF_2+1B_ complex) with half-transparent densities shown for PA (white), PA’ (light blue) and the LF (^1^LF - pink; ^2^LF - gold; ^3^LF – cyan). Notably, only ^2^LF interacts with the α-clamp.

**Figure 3.**
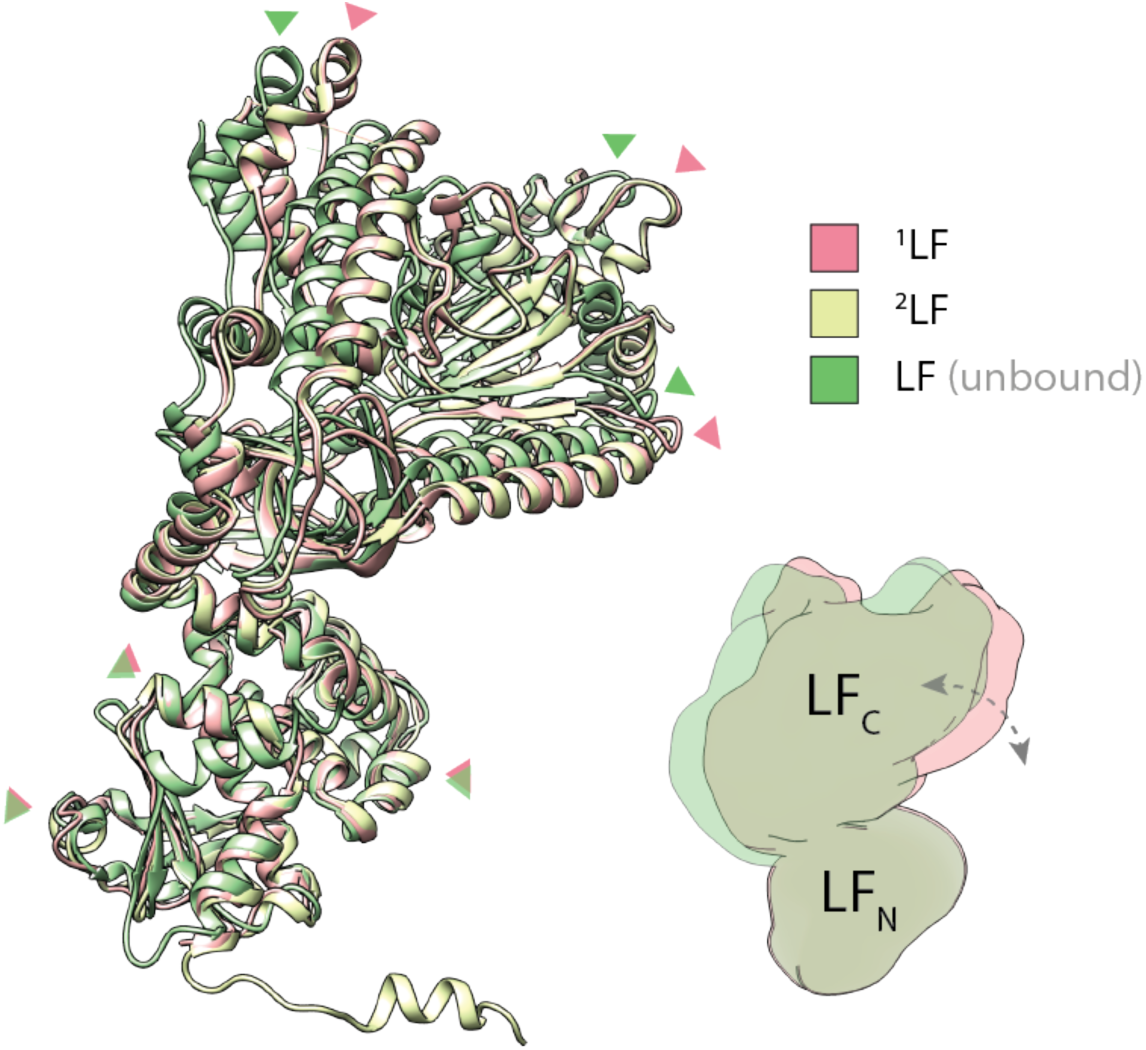
Conformational change of LF upon PA binding. Superposition of ^1^LF (green), ^2^LF (yellow) and unbound LF (green, PDB: 1J7N), aligned via their N-terminal domains. Red and green arrows indicate similar positions in ^1^LF and unbound LF, respectively. When compared with the crystal structure of the unbound lethal factor, the three LFs undergo a conformational change upon interaction with PA_7_. The C-terminal domain rotates with respect to the N-terminal domain such that the LFs come closer to form a continuous chain of head-to-tail interactions. A schematic representation illustrates the rotation of the C-terminal domain that occurs between unbound (green) and bound (red) LF conformation. See also movie S1.

**Figure 4.**
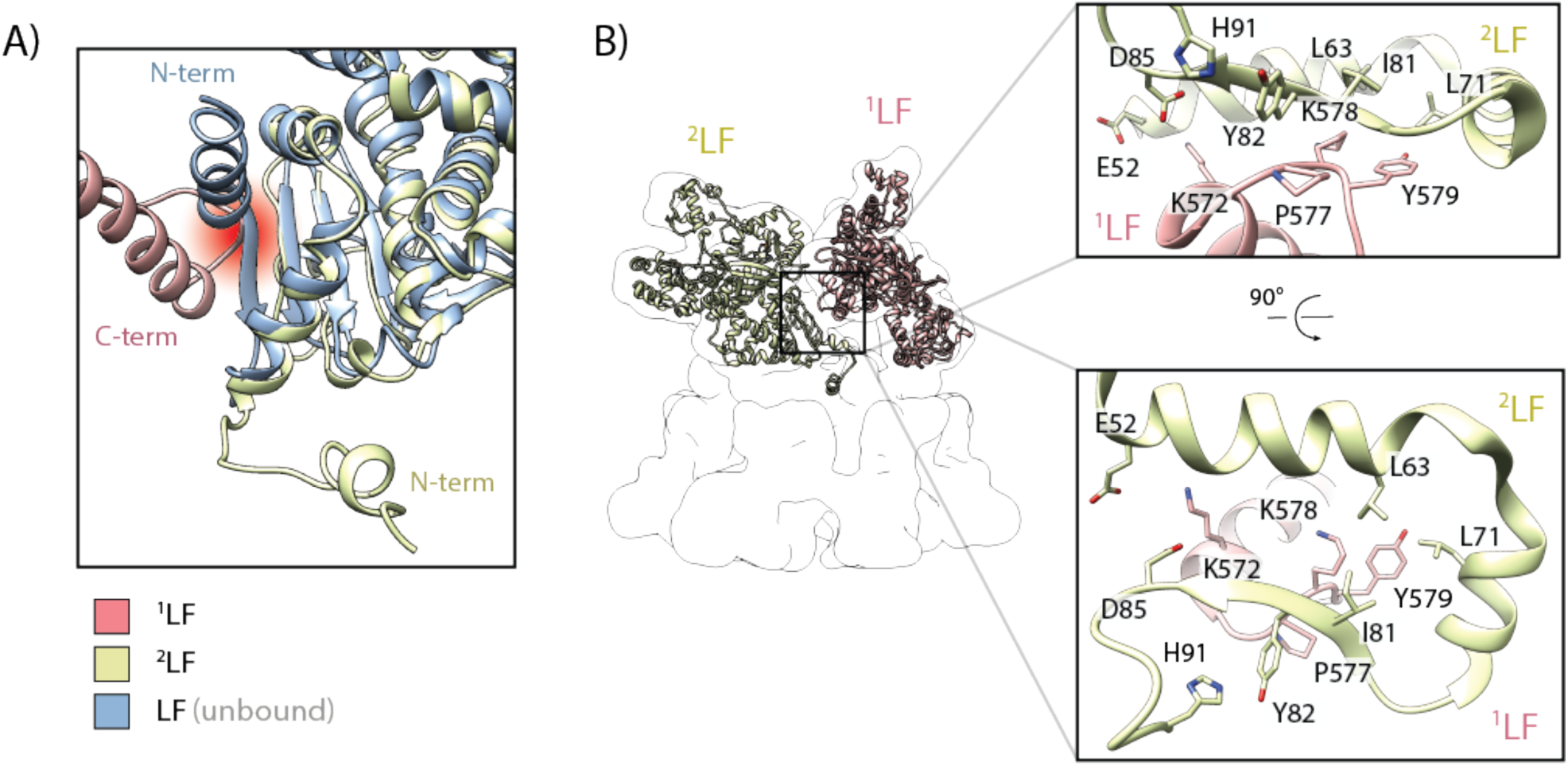
Molecular interface between two lethal factors. (**A**) Potential steric clash between the C-terminal domain of LF (red) and the N-terminal domain of an adjacent LF when it adopts the “closed” conformation (blue). The clash is highlighted as fading red spot in the background. In contrast, the “open” conformation (gold), i.e. the N-terminal α-helix interacts with the α-clamp region of PA, does not result in a steric clash. (**B**) A relatively small interaction interface mediates binding of the C-terminal domain of LF to the N-terminal domain of an adjacent LF. The positions of ^1^LF (pink) and ^2^LF (gold) are shown relative to the overall shape of the complex that is represented as transparent, low-pass filtered volume. A black square indicates the interaction interface between the two LFs. Insets show close-ups of the interacting regions in different orientations, with contributing residues labeled.

As described above, the LFs interact via their N- and C-terminal regions. In two of our structures, PA_7_LF_2+1A_ and PA_7_LF_2+1B_, two LFs only interact at one position which is located next to the major LF-PA interface. In the third structure, which we designate as PA_7_LF_(2+1A)’_, two LFs likely interact with each other also via their C-terminal region close to the central axis of the complex (Fig. 5, Movie S2). However, the position of the interaction differs from the additional interface, that has been proposed by Fabre et al. (34). At the main ^2^LF-^1^LF interface, the helix-loop region (residues 572-579) of the first lethal factor (^1^LF) forms a relatively small interface with the helix-helix-β-sheet motif (residues 52-84) of the adjacent lethal factor (^2^LF) (Fig. 4B). Residues L63, L71 and I81 of ^2^LF form a central hydrophobic cavity that interacts with Y579 of ^1^LF. In the β-sheet region of ^2^LF, we identified a potential backbone-backbone hydrogen bond between K578 and I81 of ^1^LF. In addition, P577 forms a hydrophobic interaction with Y82, which is further stabilized by H91. K572, being located on the α-helix next to the loop region in ^1^LF, could potentially form a salt bridge interaction with E52 or D85 of ^2^LF. Together these interactions mediate the binding between two LFs. Although the local resolution at the ^2^LF-^3A^LF and ^3B^LF-^1^LF interfaces does not allow the fitting of side chains (Fig. S4A-D), we could flexibly fit in the structures of ^1^LF and ^2^LF at this position. Since all structures are almost identical at backbone level (RMSD of 0.84 Å and 0.96 Å) (Fig. S5F), we expect them to exhibit a similar network of interactions. Both interfaces, LF-PA and LF-LF that we describe here limit the freedom of movement mainly in the N-terminal region of LF, but still allows a certain level of flexibility in the rest of the protein.

**Figure 5.**
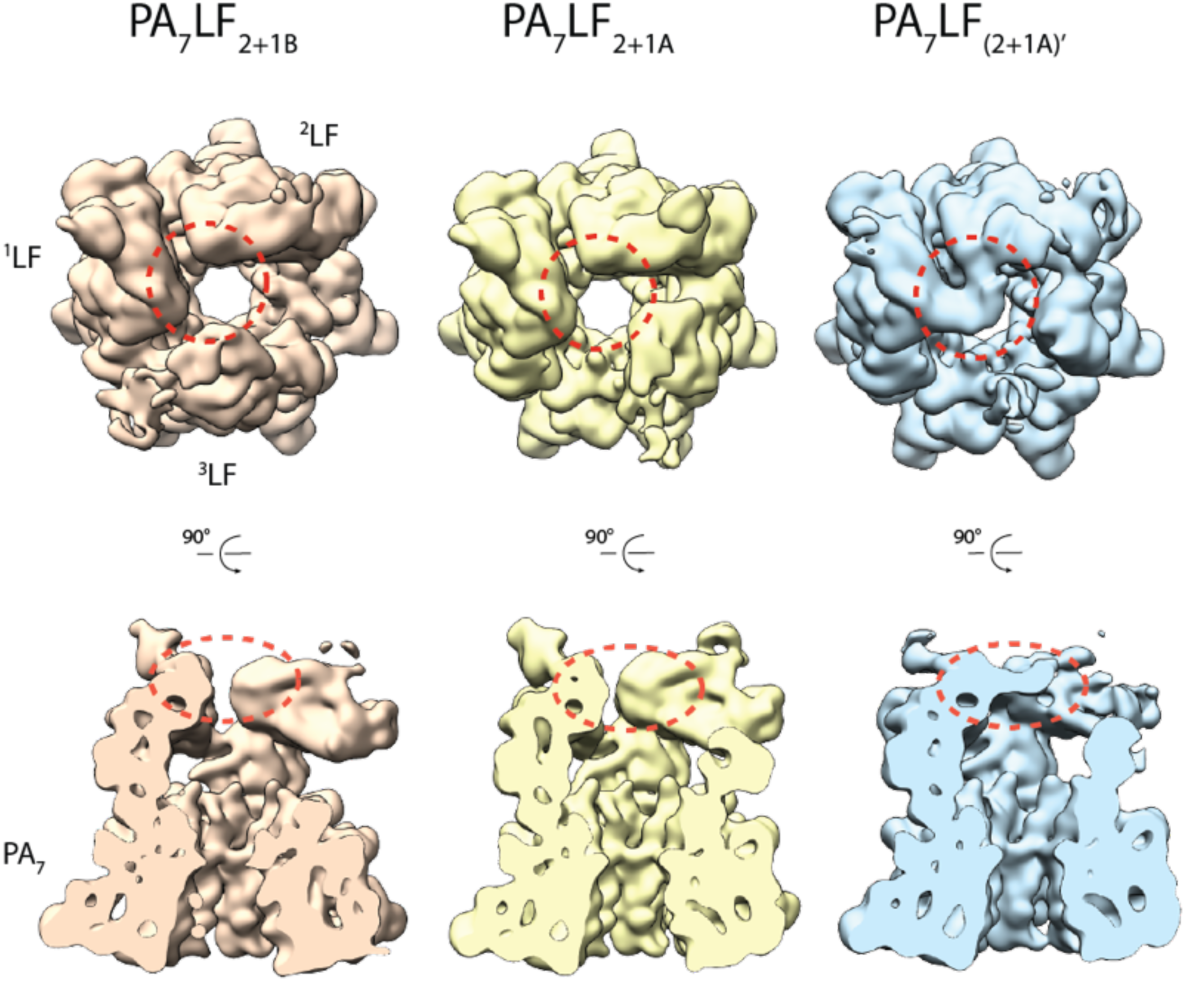
LFs can interact via their C-terminal domain. Top and side views of the low-passed filtered maps of the three PA_7_LF_3_ complexes, with PA_7_LF_2+1B_ in orange, PA_7_LF_2+1A_ in yellow and PA_7_LF_(2+1A)’_ in light blue. Volumes are shown at the same threshold. While the three LFs interact in all structures via their N- and C-terminal domains in a head-to-tail manner, an additional interface was identified in the PA_7_LF_(2+1A)’_ reconstruction. Here, the C-terminal domains of ^1^LF and ^2^LF, interact with each other close to the central axis of the PA_7_LF_3_ complex. This region is highlighted by dashed red circles. See also movie S2.

In all structures, the LFs show a gradient in flexibility (Fig. 1A-C, S4A-D). This was previously not observed at lower resolution (34). ^1^LF is resolved best in all structures, followed by ^2^LF and ^3^LF has the weakest density in all reconstructions. Since the N-terminal domain is well resolved in all LFs, this cannot be due to a varying occupancy of the binding sites, but must stem from a flexibility of the C-terminal domain. As expected, all free C-terminal domains, i.e. those that are not stabilized by an adjacent LF are more flexible than those with a binding partner. However, there is one exception, namely ^2^LF. In this case, the C-terminal domain is always flexible, independent of a stabilizing binding partner. Interestingly, ^2^LF is also the only lethal factor where the N-terminal α-helix of LF is ordered and resides in the α-clamp, suggesting that this interaction results in a destabilization of the C-terminal domain of the molecule. This is in line with a recently reported structure of the PA_7_LF_1_ pore state where the C-terminal domain of the single LF bound was not resolved while the N-terminal α-helix is also bound to the α-clamp (36).

## Discussion

We determined three structures of the fully-loaded heptameric anthrax lethal toxin complex, which differ in the position and conformations of the bound LFs. Due to a symmetry mismatch, three LFs occupy six binding sites of the heptameric PA_7_ complex, leaving one PA site empty. Compared to the “closed” state as observed in the crystal structure of LF (27), the C-terminal domain of the LFs in PA_7_LF_3_ is rotated respective to the N-terminal domain. However, only 2LF adopts the “open” conformation which was reported for the structure of PA_8_LF_4_ (21), i.e. the N-terminal α-helix interacts with the α-clamp of PA. ^1^LF and ^3^LF do not show this interaction, but can also not be in the “closed” conformation because of a steric clash with an adjacent LF. We therefore propose that they are in an “intermediate” state between toxin assembly and translocation.

Why has this state not been observed in the crystal structure of PA_8_LF_4_? It could have been missed due to averaging of the asymmetric unit of the PA_8_LF_4_ crystals, which is composed of two PAs and one LF. Another possibility is that compared to PA_7_, the PA_8_ pre-pore provides more space for the N-termini of the LFs to arrange in the “open” conformation in comparison to the PA_7_ pre-pore. However, if all LFs were indeed in a “ready-to-be-translocated position” which LF would then be translocated first through the narrow PA pore that only allows the passage of a single unfolded LF at a time? The process could in principle be stochastic, but our PA_7_LF_3_ structures offer an alternative explanation.

Already based on the low-resolution structure of the PA_7_LF_3_ pre-pore (34), it has been suggested that the order of translocation is non-stochastic and that the first LF, whose N-terminal domain is not interacting with an adjacent LF, is translocated first. After the translocation of this factor, the second LF would be released from the inhibitory bond of the first LF and then be translocated and so on (34). However, our data indicate that this chain reaction is rather unlikely.

Although we can as well only speculate about the exact order of translocation, based on our cryo-EM structures, two alternative scenarios are conceivable: In the first one, the factor in the “open” state, ^2^LF, is translocated first, followed by ^1^LF or ^3^LF which are in the “intermediate” state. The second possibility would be that ^1^LF and ^3^LF are translocated before ^2^LF. Due to the different arrangements in the complexes, both alternatives exclude a chain reaction. In addition, the translocation is not triggered or blocked by an adjacent LF.

While we cannot exclude the second scenario, we think that the first one is more likely. Being in the “open” conformation, the N-terminal α-helix of ^2^LF interacts with the α-clamp of PA. Similar to other unfolding machineries such as ClpA/Hsp100 (39), the α-clamp is known to unfold polypeptides in a sequence-independent manner. The current theory is that it first stabilizes unfolding intermediates, and introduces mechanical strain before the unfolded structure is fed further down the central pore (21). In this way it would facilitate the rapid unfolding of the entire ^2^LF molecule upon transition into the pore state. We therefore believe that ^2^LF is translocated before ^1^LF and ^3^LF. The higher flexibility in the C-terminal domain of ^2^LF in the presence of potentially stabilizing neighboring LFs suggests that the interaction of the N-terminal domain with the α-clamp results in a destabilization of the molecule. This in turn lowers the energy barrier for the unfolding of the entire LF molecule and further supports the assumption that ^2^LF is translocated first. Once ^2^LF is translocated, either ^1^LF or ^3^LF can follow. As these two LFs both adopt an “intermediate” conformation in our structures, we cannot predict which LF is translocated next.

We propose that ^2^LF is not only the first LF being translocated, but also the first one that binds to PA_7_ during toxin assembly and predict the following model (Fig. 6). Upon binding to PA, ^2^LF undergoes a conformational change from the “closed” to the “open” state (Fig. 6A, B). In the next step, ^1^LF binds to the position next to ^2^LF (Fig. 6C). Instead of transitioning into the “open” conformation, it adopts an “intermediate” conformation. The third LF binds in a similar manner, but can attach to two different PA sites, resulting in two different complexes (Fig. 6D). In this way, the assembled toxin has three LFs bound to PA_7_ with two in an “intermediate” and one in the “open” conformation (Fig. 6D).

**Figure 6.**
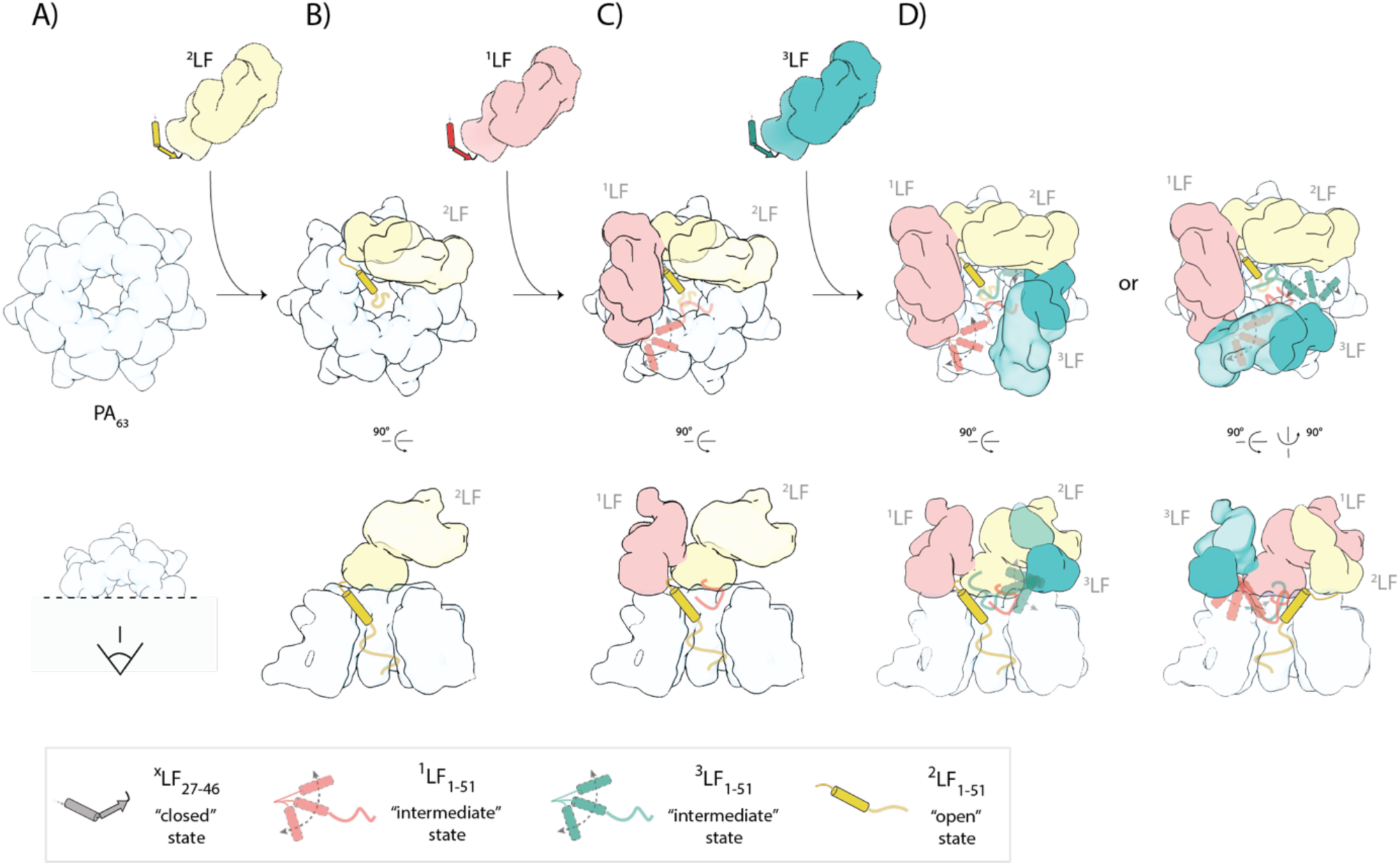
Model for PA_7_LF_3_ assembly. (**A**) After PA_7_ formation on the surface of the host cell, (**B**) ^2^LF binds to PA_7_ and its N-terminal α-helix interacts with the α-clamp region of PA, characteristic for the “open” conformation. (**C**) In the next step, ^1^LF binds adjacent to ^2^LF. Instead of undergoing a conformational change from “closed” to “open” state, it adopts an “intermediate” state. (**D**) Binding of the third LF is similar as for ^1^LF, resulting in a continuous chain of head-to-tail interactions. However, it can attach to two different PA sites, resulting in two different PA_7_LF_3_ complexes. In this way, one LF adopts the “open” conformation, whereas the other two LFs remain in an “intermediate” state.

In summary, our high-resolution cryo-EM structures provide us with novel insights into the organization of the fully-loaded heptameric anthrax lethal toxin and thus advance our understanding of toxin assembly and translocation.

## Material and Methods

### Protein expression and purification

Protective antigen (PA) from *Bacillus anthracis* was cloned into a pET19b vector (Novagen), resulting in a N-terminal His_10_-tag fusion construct. *E. coli* BL21(DE3) were transformed with the pET19b::His_10_-PA plasmid and expression was induced immediately after transformation by the addition of 75 μM IPTG. Following incubation at 28 °C for 24 h in LB medium, cells were pelleted, resuspended in lysis-buffer (20 mM Tris-HCl pH 8.5, 300 mM NaCl, 500 μM EDTA, 5 μg/ml DNAse, 1 mg/ml Lysozym plus Protease inhibitor cOmplete tablets from Sigma Aldrich) and lysed by sonication. Soluble proteins were separated from cell fragments by ultracentrifugation (15,000 rpm, 45 min, 4°C) and loaded onto Ni-IDA beads (Cube Biotech). After several washing steps, the protein was eluted with elution buffer (500 mM imidazole, 20 mM Tris-HCl pH 8.5, 500 mM NaCl, 1 mM EDTA). Protein-containing fractions were pooled and dialyzed against buffer containing 50 mM Tris-HCl pH 8.5, 150 mM NaCl, 1 mM EDTA. Subsequently, the sample was further purified using anion-exchange Mono Q (GE Healthcare) with a no-salt buffer (20 mM Tris-HCl pH 8.5) and high-salt buffer (20 mM Tris-HCl pH 8.5, 1M NaCl), applying a gradient from 0 to 40%. Next, oligomerization of PA was induced by addition of trypsin (1 μg enzyme for each mg of PA), followed by incubation on ice for 30 min. Upon addition of double molar excess of trypsin inhibitor (Sigma Aldrich), PA_7_ was further purified by size exclusion chromatography using a Superdex 200 column (GE Healthcare). Lyophilized LF (List Biological Lab. Inc., Lot#1692A1B) were resuspended in water according to the manufacturer’s manual and mixed with PA_7_ in a molar ration of 10:1. Ultimately, loaded complexes were further purified in a final size exclusion chromatography step (20 mM Tris-HCl pH 8.5, 150 mM NaCl) using a Superdex 200i column (GE Healthcare), before being used in down-stream applications.

### Reconstitution of PA_7_ in lipid-mimetic systems

For nanodisc insertion, Ni-NTA column material was first washed with ddH_2_0 and subsequently equilibrated with buffer A (50 mM NaCl, 20 mM Tris-HCl – pH 8.5, 0.05 % Octyl β-D-glucopyranoside (w/v)). In the next step, 500 µL of 0.2 µM PA_7_ in the pre-pore state was added and incubated for 25 min at room temperature. An additional washing step with buffer A was performed to remove unbound PA_7_ pre-pore, followed by a 5 min incubation step with 1 M urea at 37°C and another wash with buffer A. MSP1D1:POPC:sodium cholate ratio and preparation was done according to Akkaladevi et al (33). After dialysis (MWCO of 12-14k) for 24 to 72 h against buffer B (50 mM NaCl, 20 mM Tris-HCl pH 7.5), excess of nanodiscs was collected from five washing steps with 500 µL of buffer B. To elute PA_7_ pores inserted into nanodiscs, column material was incubated for 10 min on ice in buffer C (500 mM NaCl, 50 mM Tris-HCl pH 7.5, 50 mM imidazole). The eluted sample was concentrated and subsequently used for negative staining EM.

For the preparation of pre-formed liposomes, POPC was initially solubilized in 5 % OG. Solubilized lipids were dialyzed (MWCO: 12-14k) for 8 - 12 h at 4°C against buffer A and subsequently PA_7_ pre-pores were added to the lipids in a 1 : 10 molar ratio. Following 24 – 72 h dialysis (MWCO:12-14k) against buffer D (50 mM NaCl 50 mM NaOAc, pH 5.0), samples were used for negative staining EM.

### Negative-stain electron microscopy

Complex purity and integrity were assessed by negative stain electron microscopy prior to cryo-EM grid preparation and image acquisition. For negative stain, 4 μl of purified PA_7_LF_3_ complex at a concentration of ∼0.04 mg/ml was applied onto a freshly glow discharged carbon-coated copper grid (Agar Scientific; G400C) and incubated for 45 s. Subsequently, excess liquid was blotted away with Whatman no. 4 filter papers. The sample was stained with 0.8 % (w/v) uranyl acetate (Sigma Aldrich). Micrographs were recorded manually using a JEOL JEM-1400 TEM, operated at an acceleration voltage of 120 kV, equipped with a 4,000 × 4,000 CMOS detector F416 (TVIPS) and a pixel size of 1.84 Å/px.

### Sample vitrification

For Cryo-EM sample preparation, 4 μl of purified PA_7_LF_3_ at a concentration of ∼0.06 mg/ml was applied onto freshly glow discharged grids (Quantifoil R 1.2/1.3 holey carbon with a 2 nm additional carbon support) and incubated for 45 s. Subsequently, grids were blotted automatically and plunged into liquid ethane using a CryoPlunge3 (Gatan) at a humidity of ∼ 95 %. Grid quality was screened before data collection using a JEOL JEM-1400 TEM electron microscope (same settings as for negative-stain electron microscopy) or with an Arctica microscope (FEI), operated at 200 kV. Grids were kept in liquid nitrogen for long-term storage.

### Cryo-EM data acquisition

Cryo-EM data sets of PA_7_LF_3_ were collected on a Titan Krios transmission electron microscope (FEI) equipped with a high-brightened field-emission gun (XFEG), operated at an acceleration voltage of 300 kV. Micrographs were recorded on a K2 direct electron detector (Gatan) at 130,000 x magnification in counting mode, corresponding to a pixel size of 1.07 Å. 40 frames taken at intervals of 375 ms (1.86 e^-^/Å^2^) were collected during each exposure, resulting in a total exposure time of 15 s and total electron dose of 74.4 e^-^/Å^2^. Using the automated data collection software EPU (FEI), a total of 5238 micrographs with a defocus range between -1.2 and -2.6 µm was automatically collected.

### Image processing and 3-D reconstruction

Micrographs of the dataset were inspected visually and ones with extensive ice contamination or high drift were discarded. Next, frames were aligned and summed using MotionCor2 (in 3 x 3 patch mode) (40). By doing so, dose-weighted and un-weighted full-dose images were generated. Image and data processing were performed with the SPHIRE/EMAN2 software package (41). Un-weighted full-dose images were used for defocus and astigmatism estimation by CTER. With the help of the drift assessment tool in SPHIRE, drift-corrected micrographs were further sorted to discard high defocus as well as high drift images that could not be compensated for by frame alignment.

For the PA_7_LF_3_ dataset, particles were automatically selected based on a trained model using the crYOLO software, implemented in SPHIRE (42). In total, 382 k particles were extracted from the dose-weighted full dose images with a final window size of 336 x 336 pixel. Two-dimensional classification was performed using the iterative and stable alignment and clustering (ISAC) algorithm implemented in SPHIRE. Several rounds of 2-D classification yielded a total number of 213 k ‘clean’ dose-weighted and drift-corrected particles. During the manual inspection of the 2-D class averages, top views of the particles were excluded.

A generated composite crystal structure consisting of PA_7_ (PDB:1TZO) decorated with three full-length LF (PDB:1J7N), docked with their N-terminal domain to PA as observed in the PA_8_LF_4_ structure (PDB: 3KWV), was converted into electron density (sp_pdb2em functionality in SPHIRE). After filtering to 30 Å, this map served as reference in the subsequent 3-D refinement. The 3-D refinement without imposed symmetry (sxmeridien in SPHIRE, C1) yielded an initial 3.5 Å electron density map of the PA_7_LF_3_ complex. Several rounds of 3-D classification and rotation of certain classes were necessary to separate particles belonging to PA_7_LF_2+1A_, PA_7_LF_2+1B_ and PA_7_LF_(2+1A)’_ complexes. The flowchart of the image processing strategy including the obtained 3-D classes as well as the number of particles that they contained is described in detail in Fig. S3.

Global resolutions of the final maps were calculated between two independently refined half maps at the 0.143 FSC criterion, local resolution was calculated using sp_locres in SPHIRE. The final densities were filtered according to local resolution or the local de-noising filter LAFTER was applied to recover features with more signal than noise (based on half maps) (43).

### Model building, refinement and validation

To build the PA_7_ model, a single monomer of the PA_7_ crystal structure (PDB:1TZO) was used as starting model and a preliminary fit into the PA density of the PA_7_LF_3-masked_ map was done using rigid body fitting in Chimera. Next, it was flexibly fitted into the corresponding density using iMODFIT (44). The resulting model was copied and fitted to the other six PA densities and each monomer was separately refined further using a combination of manual model building in COOT and real-space refinement in PHENIX. Subsequently, all seven monomers were merged together to create the final model of PA_7_. Unresolved loop regions were deleted (275-285, 301-322, 424-428 and 644-656) and less resolved regions exchanged to poly-A (641, 666-700, 710-715 and 720-735).

For the model building of the lethal factors, a composite model of residues 29 to 250 from the N-terminal domain of LF (PDB:3KWV) and residues 251 to 773 from the full-length LF structure (PDB:1J7N) was generated. This hybrid pdb served as staring model and was initially fitted into the density of ^1^LF and ^2^LF in the PA_7_LF_3-masked_ structure using rigid body fitting in Chimera. In the next step, models were flexibly fitted into the density using iMODFIT, followed by further refinement using a combination of manual model building in COOT and real-space refinement in PHENIX for ^1^LF (52-254 and 550-600) and ^2^LF (32-253).

The resulting models for ^1^LF, ^2^LF and PA_7_ served again as starting point for the PA_7_LF_2+1B_ structure and were flexibly fitted into the corresponding density. Additional refinement using a combination of manual model building in COOT and real-space refinement in PHENIX was performed for ^1^LF (52-254 and 550-600) and ^2^LF (32-253), similar as in the PA_7_LF_3-masked_ structure. The density for the N-terminal domain of ^3^LF was less well resolved and therefore only flexibly fitted into the density using iMODFIT (52-254), whereas the C-terminal domain was fitted using the ‘rigid body fit’ tool in Chimera.

Like for the PA_7_LF_2+1B_ structure, obtained models of ^1^LF, ^2^LF, ^3^LF (of the PA_7_LF_2+1B_ structure) and PA_7_, were flexibility fitted into the PA_7_LF_2+1A_ density map to obtain the final model of PA_7_LF_2+1A_. The C-terminal domain of ^3^LF was fitted using the ‘rigid body fit’ tool in Chimera. Geometries of the final refined models were obtained from PHENIX with data statistics summarized in Table S1.

### Structure analysis and visualization

UCSF Chimera was used for structure analysis, visualization and figure preparation. The angular distribution plots as well as beautified 2-D class averages were calculated using SPHIRE.

## Acknowledgments

We thank O. Hofnagel and D. Prumbaum for assistance with data collection. This work was supported by the Max Planck Society (to S.R.) and the European Council under the European Union’s Seventh Framework Programme (FP7/ 2007–2013) (grant no. 615984) (to S.R.). D.Q. is a fellow of Fonds der Chemischen Industrie.

## Author contributions

S.R. designed the project. K.A. and A.E.L. provided protein complexes. C.A. prepared specimens, recorded and processed the EM data. C.A., D.Q. and C.G. analyzed the data. D.Q. prepared figures. S.R. managed the project. D.Q. and S.R. wrote the manuscript with input from all authors.

## Data availability

The cryo-EM density maps of the PA_7_LF_2+1A_, PA_7_LF_2+1B_ and PA_7_LF_(2+1A)’_, complexes are deposited into the Electron Microscopy Data Bank with the accession codes EMD-xxxx, EMD-xxxx and EMD-xxxx, respectively. Corresponding coordinates for PA_7_LF_2+1A_ and PA_7_LF_2+1B_ have been deposited in the Protein Data Bank under accession number xxxx and xxxx. Relevant data and details of plasmids and strains are available from the corresponding author upon reasonable request.

## Supporting information figure captions

**Figure S1.**
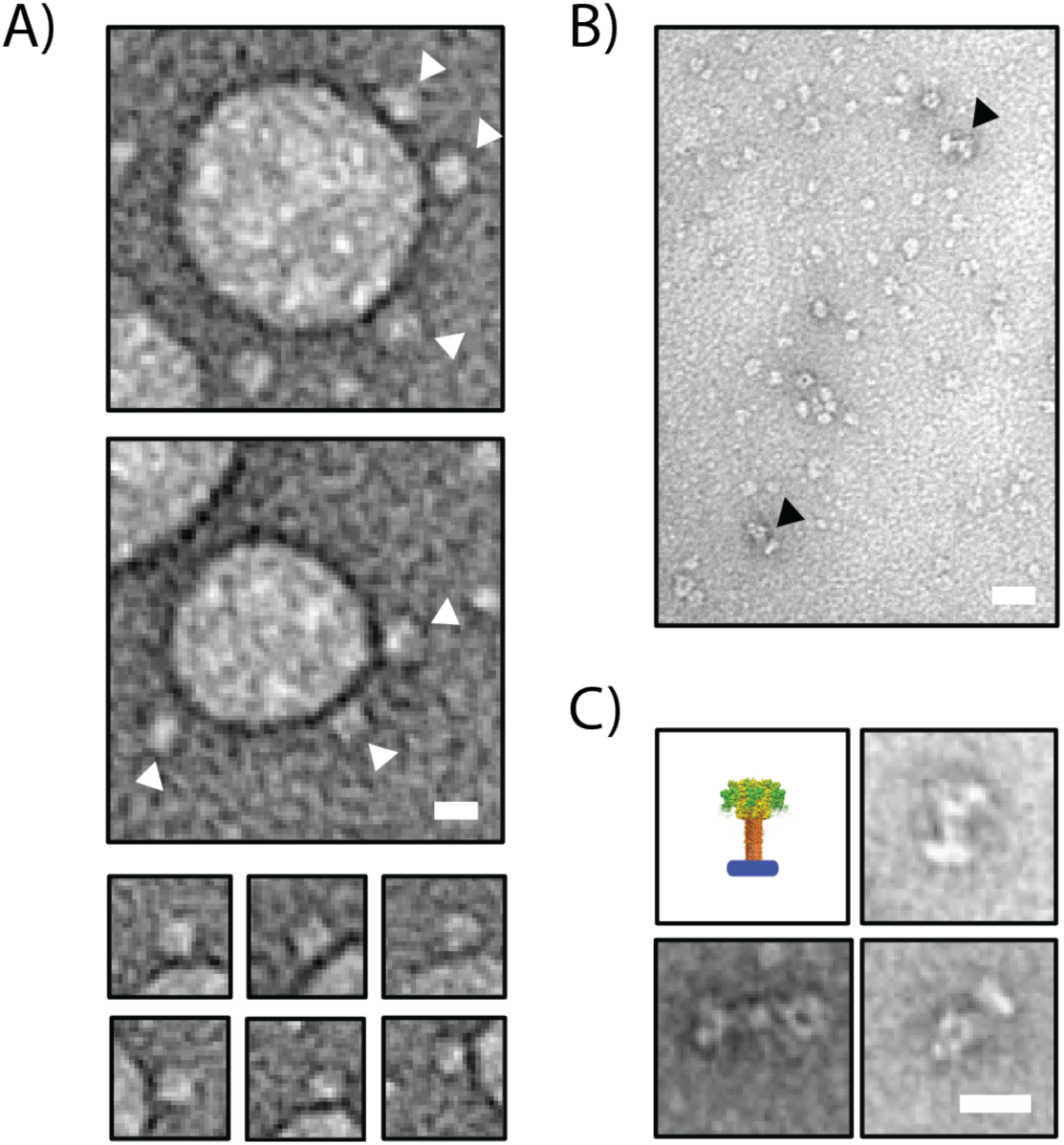
Reconstitution of PA_7_ into lipid mimetic systems after pore transition. (**A**) Representative negatively stained electron micrograph areas of PA_7_ reconstituted into POPC liposomes (top panels), with individual inserted particles highlighted by white arrowheads. Selection of inserted particles in smaller lipid vesicles (bottom panel). Scale bar: 15 nm. Particles are clearly accumulated at lipid membranes. (**B**) Representative negatively stained electron micrograph area of PA_7_ reconstituted in lipid nanodiscs (MSP1D1), with individual inserted particles highlighted by black arrowheads. Scale bar: 20 nm (**C**) Model of PA_7_ complexes inserted into lipid nanodiscs with additional examples of individual particles after reconstitution (same nanodiscs as in B). Scale bar: 20 nm.

**Figure S2.**
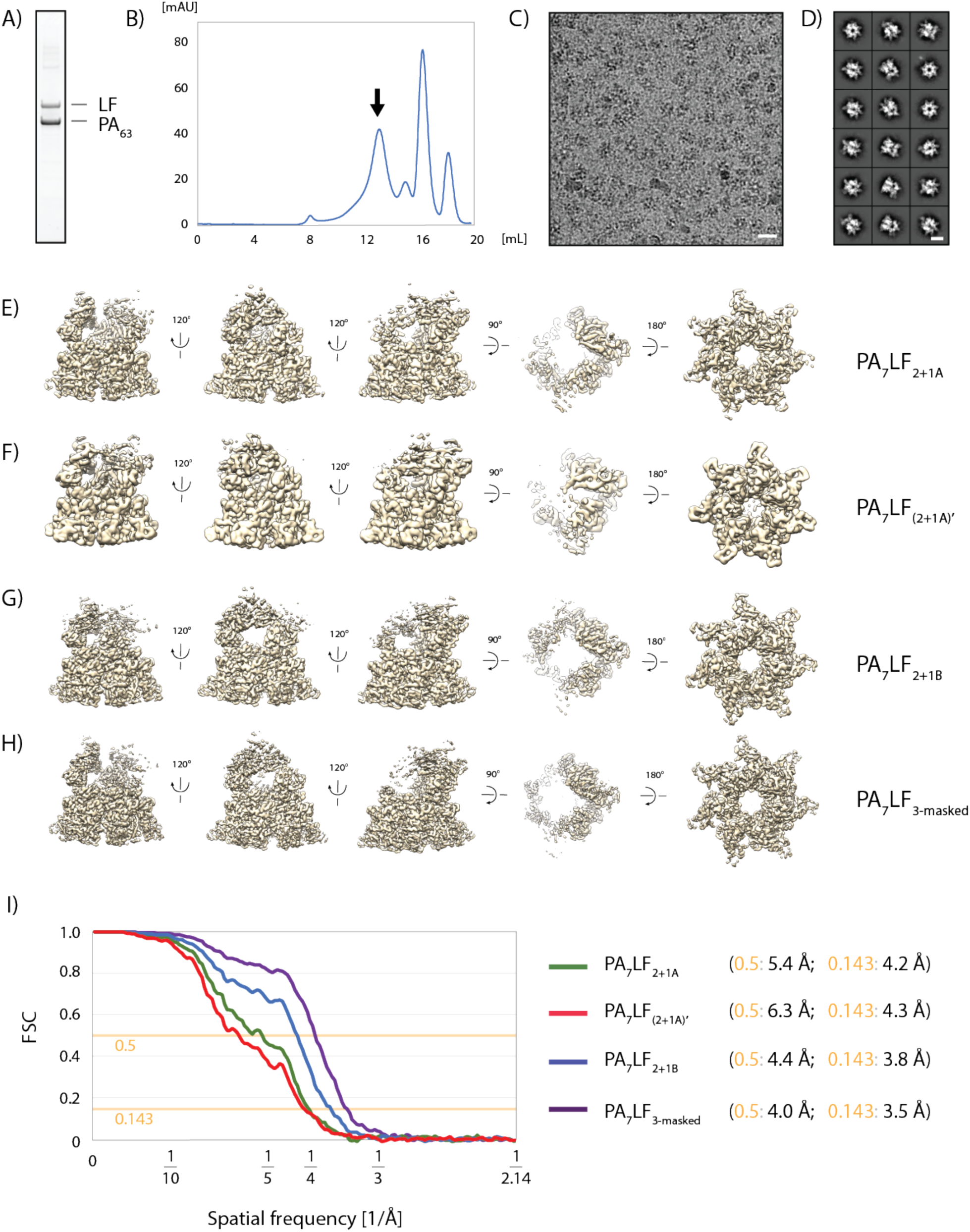
Purification and cryo-EM of PA_7_LF_3_. (**A**) Coomassie-stained SDS-PAGE of purified PA_7_LF_3_ complex. (**B**) Size exclusion chromatography profile of the PA_7_LF_3_ complex using a Superdex 200 column. Sample fraction used for cryo-EM studies is indicated by black arrow. (**C**) Representative digital micrograph area of vitrified PA_7_LF_3_ complex. Scale bar: 20 nm. (**D**) Representative 2-D class averages corresponding to **C**. Scale bar: 10 nm. (**E-H**) Rotated views of the 3-D reconstruction of PA_7_LF_2+1A_ (**E**), PA_7_LF_(2+1A)’_ (**F**), PA_7_LF_2+1B_ (**G**), and PA_7_LF_3-masked_ (**H**), respectively. (**I**) FSC curves between two independently refined half-maps of PA_7_LF_2+1A_ (green), PA_7_LF_(2+1A)’_ (red), PA_7_LF_2+1B_ (blue) and PA_7_LF_3-masked_ (purple).

**Figure S3.**
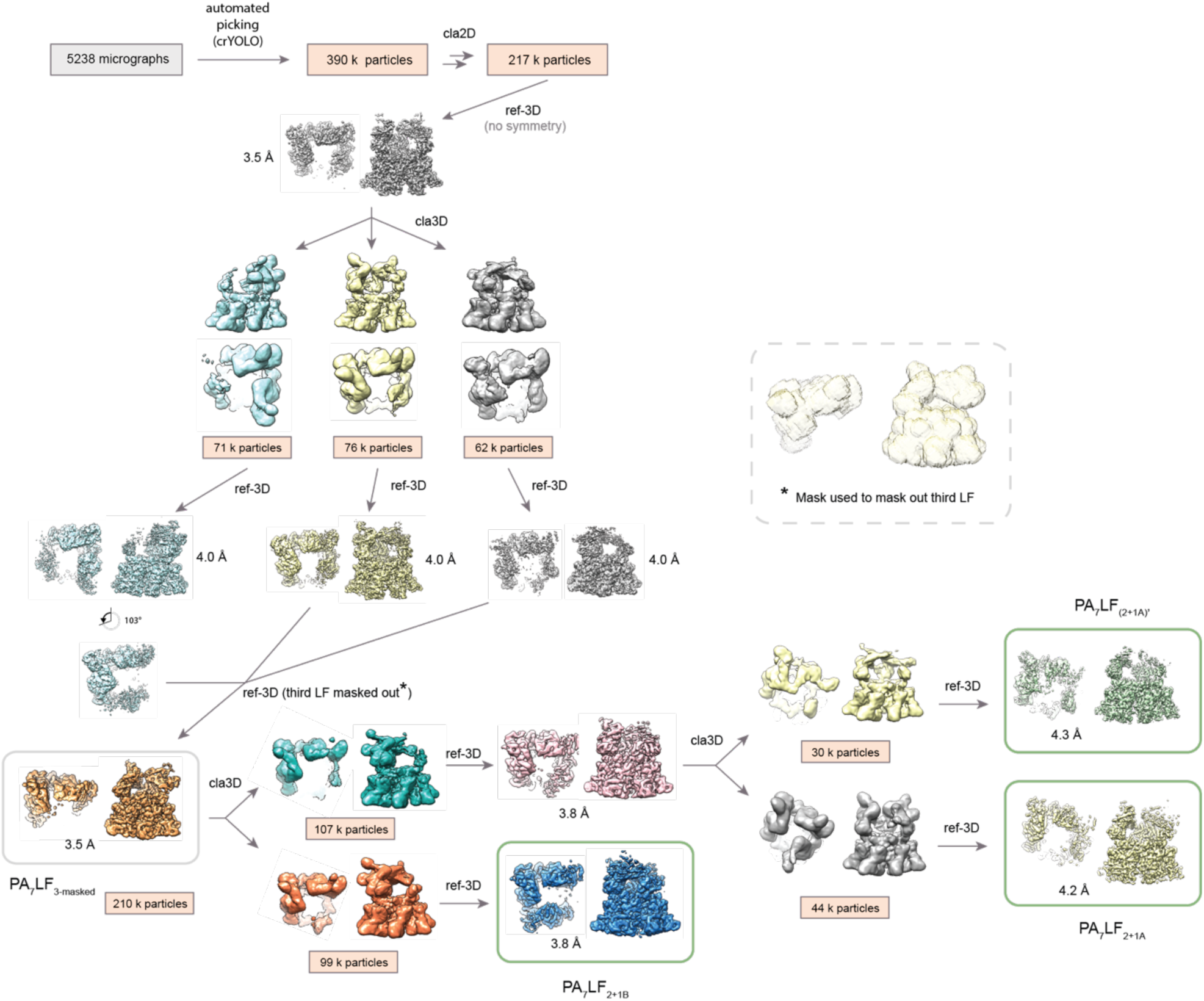
Flowchart of image processing strategy in SPHIRE. Te single particle processing workflow is shown that included multiple 3-D classification steps as well as rotation of individual classes (indicated by rotation symbol). Number of particles in each class is provided as orange box below the respective structure and the obtained resolution of the map after 3-D refinement is indicated. For each structure a top and side view is shown (in top views PA_7_ density is partially clipped to focus on the bound LFs). Mask for masking out third LF is provided in dashed box. Final electron density maps are highlighted by green boxes. Abbreviations: cla3D – 3-D classification, cla2D – 2-D classification, ref-3D – 3-D refinement.

**Figure S4.**
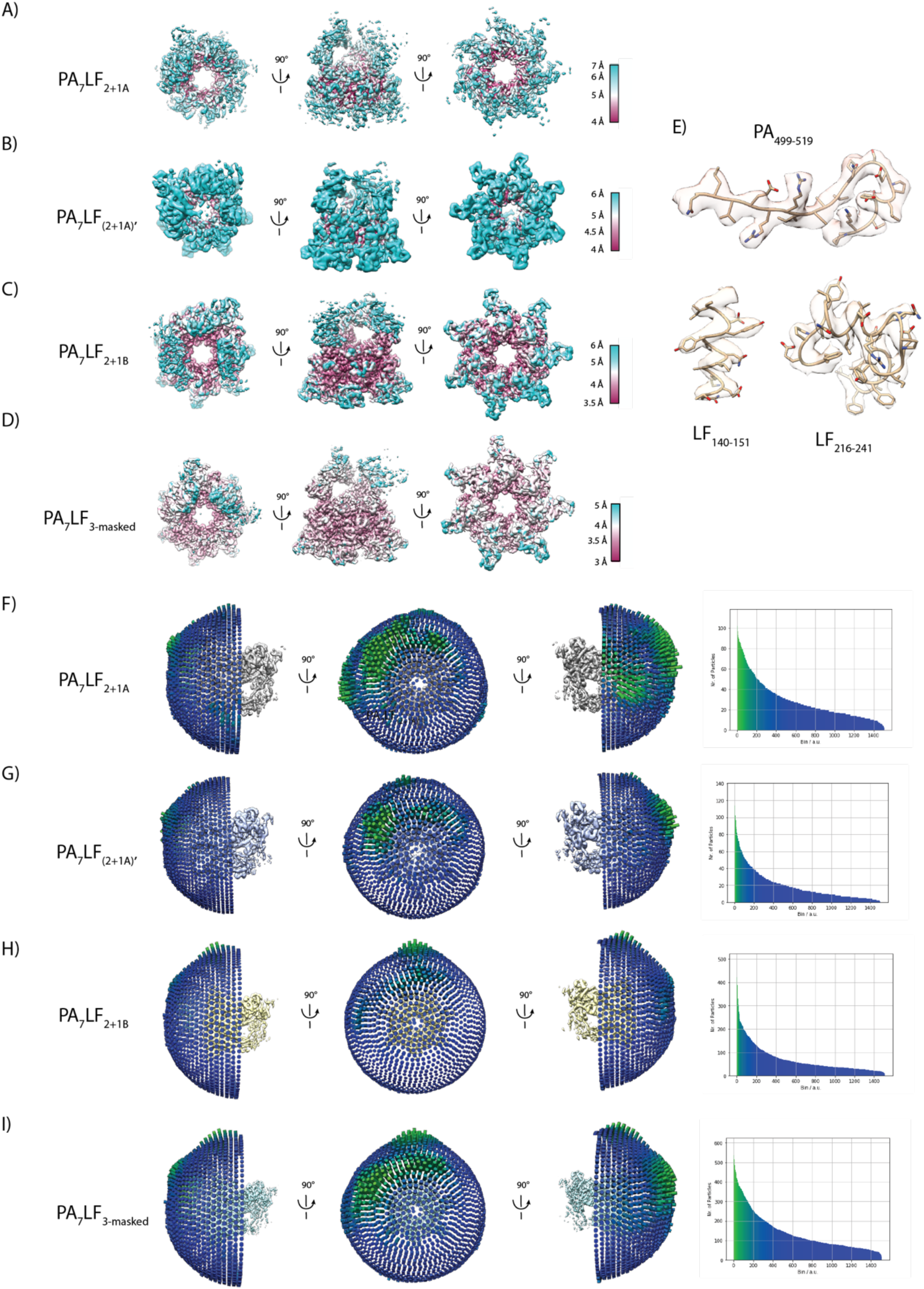
Local resolution and 3-D orientation plots. (**A-D**) Rotated views of the reconstructions, PA_7_LF_2+1A_ (**A**), PA_7_LF_(2+1A)’_ (**B**), PA_7_LF_2+1B_ (**C**), and PA_7_LF_3-masked_ (**D**), respectively, colored by local resolution. Corresponding color key of local resolution is provided on the right. (**E**) Selected examples of side chain densities corresponding to PA and LF with atomic models fitted. (**F**) Rotated views of the 3-D angular distribution plot for the PA_7_LF_2+1A_ reconstruction, in which the relative height of bars represents the number of containing particles. Corresponding 2-D histogram is shown on the right. (**G-I**) Same as in **F** for PA_7_LF_(2+1A)’_ (**G**), PA_7_LF_2+1B_ (**H**), and PA_7_LF_3-masked_ (**I**).

**Figure S5.**
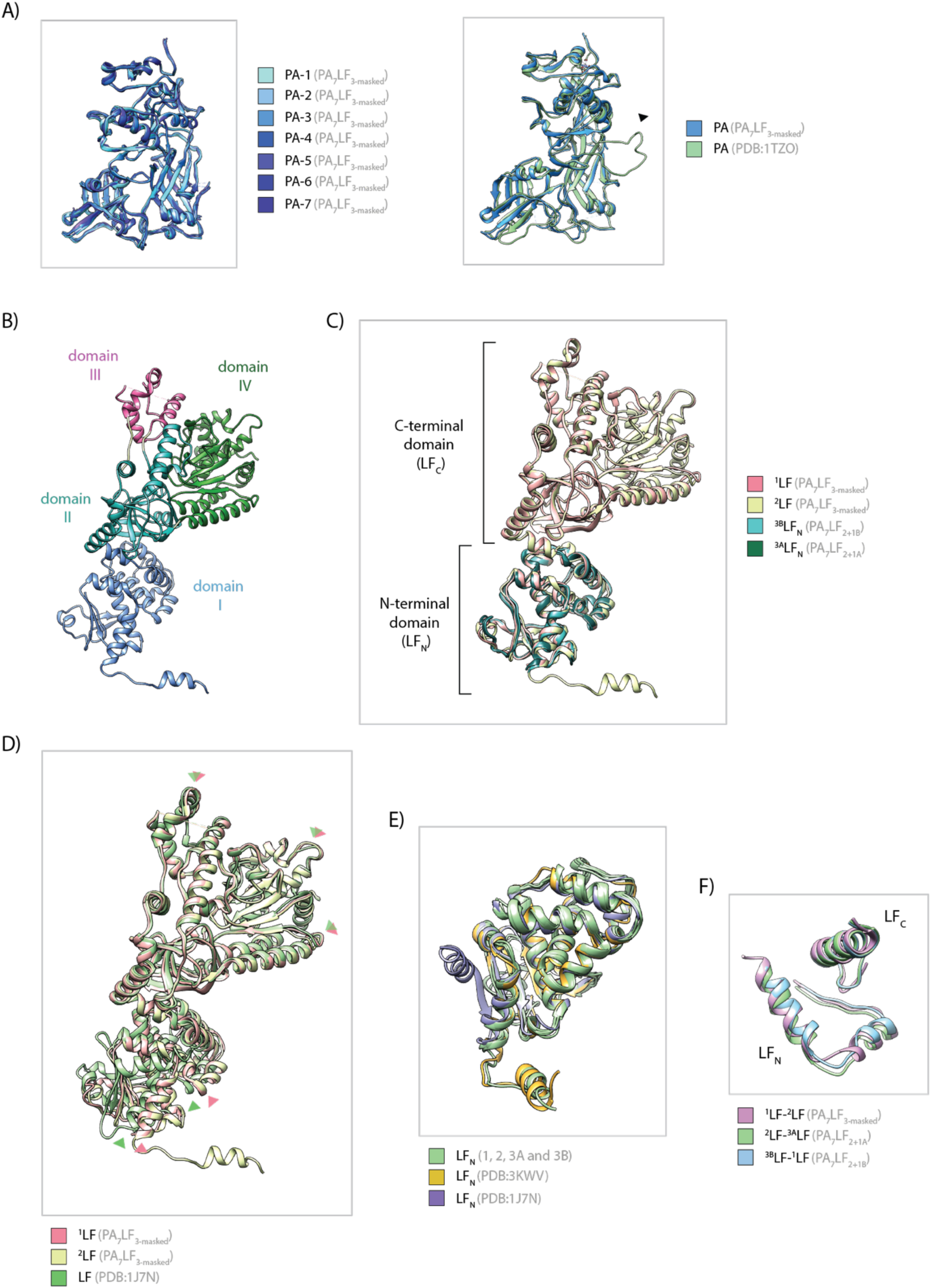
Structure comparison of PAs and LFs. (**A**) Superposition of the seven PA protomers in PA_7_LF_3_, which are colored in different blue hues (left panel), and a single PA subunit (blue) with the known crystal structure (PDB: 1TZO, green, right panel). Loop region 2β2-2β3 (residues 300-323), resolved only in the crystal structure, is highlighted by a black arrowhead. (**B**) Domain organization of LF with individual domains highlighted by different colors. (**C**) Superposition of individual LFs in the PA_7_LF_3_ structures with ^1^LF in pink, ^2^LF in gold, ^3B^LF_N_ in cyan and ^3A^LF_N_ in dark green. (**D**) Superposition of ^1^LF (pink), ^2^LF (gold) and unbound LF (PDB: 1J7N, green), aligned via their C-terminal domain. Green and red arrows indicate similar positions in ^1^LF and unbound LF (PDB:1J7N), respectively. Comparison reveals that the C-terminal domain is rotated respective to the N-terminal domain in the PA_7_LF_3_ structures. (**E**) Superposition of the N-terminal domain of the three LFs in PA_7_LF_3_ (green), of LF in the “open” conformation in PA_8_LF_4_ (PDB: 3KWV, dark yellow) and of unbound LF in the “closed” conformation (PDB: 1J7N, purple). (**F**) Superposition of the three LF-LF interfaces with ^1^LF-^2^LF in pink, ^2^LF-^3A^LF in green and ^3B^LF- ^1^LF in blue.

## Supporting information movie captions

**Movie S1. Conformational change of LF upon PA binding**.

The C-terminal domain of the three LF molecules rotate respective to the N-terminal domain upon binding to PA_7_ when compared with the unbound LF structure (PDB:1J7N), to form a continuous chain of head-to-tail interactions. Top view of the morph between both conformations is shown, with LFs in blue and PA_7_ in transparent grey.

**Movie S2. LFs can interact via their C-terminal domains**.

In our PA_7_LF_(2+1A)’_ reconstruction, two LF molecules interact with each other via their C-terminal domain close to the central axis of the complex, thus forming an additional LF-LF interface. Top view of the morph between this conformation (light blue) and the one observed in the PA_7_LF_2+1A_ (yellow) is shown. Volumes are low-pass filtered and shown at the same threshold.

## Supporting information tables

**Table S1.**
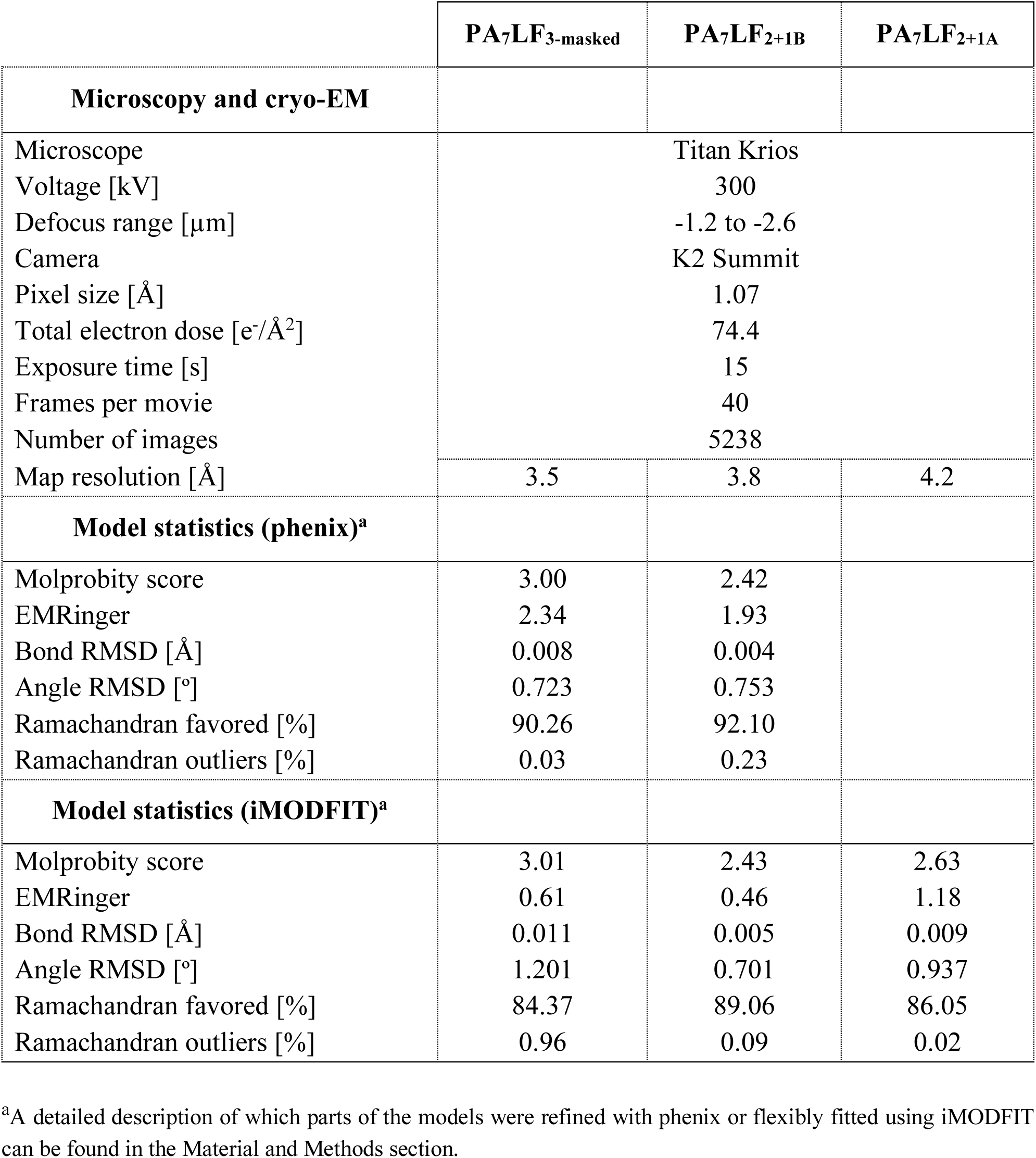
Data collection, refinement and model building statistics.

